# Retinal neuroanatomy of two emerging model organisms, the spiny mouse (*Acomys dimidiatus*) and the Mongolian gerbil (*Meriones unguiculatus*)

**DOI:** 10.1101/2024.01.17.576137

**Authors:** Jessica D. Bills, Ashley W. Seifert, Ann C. Morris

## Abstract

**Purpose:** Degenerative eye diseases such as macular degeneration and retinitis pigmentosa slowly deteriorate vision, ultimately leading to blindness. Current research with laboratory animal models largely utilizes small mammals that are nocturnal and lack the ability to restore lost vision. In contrast, the Mongolian gerbil is a diurnal rodent with good photopic vision, and the eastern spiny mouse is a small desert-dwelling rodent with remarkable regenerative capabilities. The goal of this study was to identify several antibodies that detect retinal cell classes in *Meriones* and *Acomys*, and to describe the retinal anatomy of these two species in comparison to outbred laboratory mice (*Mus musculus*).

**Methods:** Retinal sections were obtained from adult eyes and subjected to histological or immuno-staining with antibodies for various retinal cell types. Sections were imaged by light, fluorescence, and confocal microscopy, assessing cell number and morphology. Cell density, morphology, and placement were compared between species qualitatively and quantitatively.

**Results:** Immunohistochemical visualization and analysis of all general classes of retinal neurons and Müller glia revealed a classic assembly of retinal cells with a few deviations compared to *Mus*. *Meriones* displayed the highest density of cone photoreceptors and *Acomys* the lowest. A higher density of bipolar cell bodies in the proximal portion of the inner nuclear layer was observed in both *Acomys* and *Meriones* compared to *Mus*, and both species exhibited an increase in amacrine cell density compared to *Mus*.

**Conclusion:** We have characterized similarities and differences in the retinal anatomy and cellular density between *Meriones*, *Acomys*, and *Mus*. We identified several commercially available antibodies that reliably detect retinal cell types in the *Acomys* and *Meriones* retina. Our results provide a foundation for future research into the visual system adaptations of both of these interesting rodent species.

## INTRODUCTION

The retina carries out the first critical steps of light perception. The vertebrate retina contains five types of neurons (ganglion cells, amacrine cells, bipolar cells, horizontal cells, and photoreceptors), and one intrinsic glial cell type (Müller glia), organized into three anatomical layers, the ganglion cell layer (GCL), the inner nuclear layer (INL) and the outer nuclear layer (ONL). While the overall organization and general cell types found in the retina are well conserved across vertebrates, the subtype, number, and positioning of the retinal neurons can differ substantially ^1^.These differences can result in varying functional alterations that reflect adaptations to different environments and lifestyles, such as nocturnal or diurnal activity ^2–4^. Differences in visual acuity reflect variations in the organism’s light detecting neurons, the photoreceptors.

There are two general classes of vertebrate photoreceptors, rods and cones. Rod photoreceptors mediate dim-light vision and are sensitive enough to detect a single photon of light, whereas cone photoreceptors mediate daytime vision and allow for light detection across the color spectra. Most mammals have one rod type and two to three cone subtypes ^5,6^. Large differences exist in visual capabilities and photoreceptor complement across mammals. *Mus musculus*, the common house mouse, has two cone subtypes and a rod-dominant retina (a ratio of 35:1 rods to cones) which is adapted for nocturnal vision ^7,8^. In contrast, the diurnal thirteen-lined ground squirrel retina has the same cone subtypes as mouse, but with a ratio of 50:1 cones to rods ^9,10^

The Mongolian gerbil (*Meriones unguiculatus*) is a small rodent naturally inhabiting grasslands, shrublands, and deserts in China, Mongolia, and Russia ^11^. As a laboratory model, they have been utilized for research in a variety of areas, including reproduction, diabetes, epilepsy, and aging ^12–14^. As diurnal rodents, they have higher visual acuity and enhanced photopic vision compared to mice and rats ^15–17^. Previous studies describing *Meriones* retinal anatomy documented a cone-dominant retina with similar retinal layers to other vertebrates ^18–21^. However, a comprehensive assessment of retinal organization and retinal cell types is currently lacking for this species.

Spiny mice (genus *Acomys*) include over 15 species and are small desert rodents residing throughout Africa, the Middle East, and Southwest Asia. Despite the use of “mouse” in their common name, genomic sequencing shows that Deomyinea (subfamily of *Acomys)* are more closely related to Gerbillinea than to Muridae ^22^. For over four decades, these animals have been researched for their desert adapted physiology and gregarious social behavior ^23–27^. Importantly, spiny mice are precocial; they are born mobile, with open eyes, and are curious and social as soon as one day after birth ^23^.

Several *Acomys* species possess remarkable regenerative ability among musculoskeletal tissues and skin ^28–32^ and more recently it was discovered that they can completely recover from complete spinal cord transection^30,32–37^. Although there has been some published work investigating spiny mouse visual behavior, information about their retinal anatomy is limited ^38,39^. In the rocky deserts in southern Israel, *Acomys cahirinus* and a closely related sister species, *Acomys russatus*, occupy the same habitat. Previous studies have shown that *A. russatus* can switch its behavior from nocturnal to diurnal for better foraging and reduced competition^38,39^. Furthermore, a recent study examining sleeping patterns in *Acomys* compared to the strictly nocturnal *Mus* revealed that *Acomys* sleep with their eyes completely open and are active both during the day and night, with sleep primarily occurring at dusk ^40^. Given these uncommon characteristics, a comprehensive assessment of their retinal anatomy could provide insight into whether their visual adaptations correlate with abundance of specific retinal cell types and constitutes a necessary first step in investigating their retinal regenerative ability. Here we provide the first comprehensive examination of retinal anatomy in *A. dimidiatus* (*Acomys*) and *M. unguiculatus* (*Meriones*) compared to the well characterized laboratory mouse *M. musculus* (*Mus*).

## METHODS

### Animal Care

Spiny mice (*Acomys dimidiatus;* referred to as *Acomys cahirinus* in some previous reports), Mongolian gerbil (*Meriones unguiculatus)* and outbred laboratory mice (*Mus musculus*; ND4, Envigo) were housed in our animal housing facility at the University of Kentucky, Lexington, KY. All species were maintained on 14% mouse chow (Teklad Global 2014, Harlan Laboratories); spiny mice also received sunflower seeds. Animals were exposed to natural light through large windows (approximately 12L:12D light cycle). Both males and females of breeding age (6-12 months old) were utilized for all experiments. All procedures were carried out in accordance with the guidelines established by the University of Kentucky Institutional Animal Care and Use Committee (2019-3254).

### Tissue collection, histology, and immunohistochemistry

Animals were euthanized using a mixture of isoflurane gas and oxygen and eyes were enucleated with the cornea removed. Eye cups, including the retina, were fixed with 4% paraformaldehyde for immunohistochemistry and 10% neutral buffered formalin for histology. Histological samples were processed and embedded in paraffin (Leica Biosystems, Buffalo Grove, IL) and 7 μm sections were placed onto Superfrost Plus slides (Fisher Scientific). Tissue sections were processed for routine histology and stained with hematoxylin and eosin reagents (Sigma-Aldrich St. Louis, MO). For immunohistochemistry, tissues were prepared and cryoprotected by submersion in 10% followed by 30% sucrose, then embedded in Optimal Cutting Temperature compound (Sakura Finetek, CA), frozen, and sectioned at 10μm thickness. Sections were utilized for immunohistochemical processing as follows: cryosections were washed extensively in PBS and treated with citrate buffer (10 mM; pH 6.0) for 20 minutes at 96°C to improve antigen retrieval. Sections were then blocked for a minimum of 60 minutes with PBS/Tween 20 or PBS 0.1M 0.3% Triton X-100 plus 5% serum and incubated overnight with the primary antibody, diluted in blocking reagent. A list of antibodies and dilutions can be found in **Table 1**. A minimum of 16 hours later, sections were washed with PBS/Tween-20, exposed for 60 minutes to appropriate secondary antibodies (diluted in blocking reagent) away from light, washed again with PBS and then counter-stained with nuclear Hoechst (Invitrogen Waltham, MA). Slides were mounted with a 60% glycerol solution and imaged. Control labeling was performed without primary antibodies. All steps were carried out at room temperature unless otherwise noted.

**Table 1:**
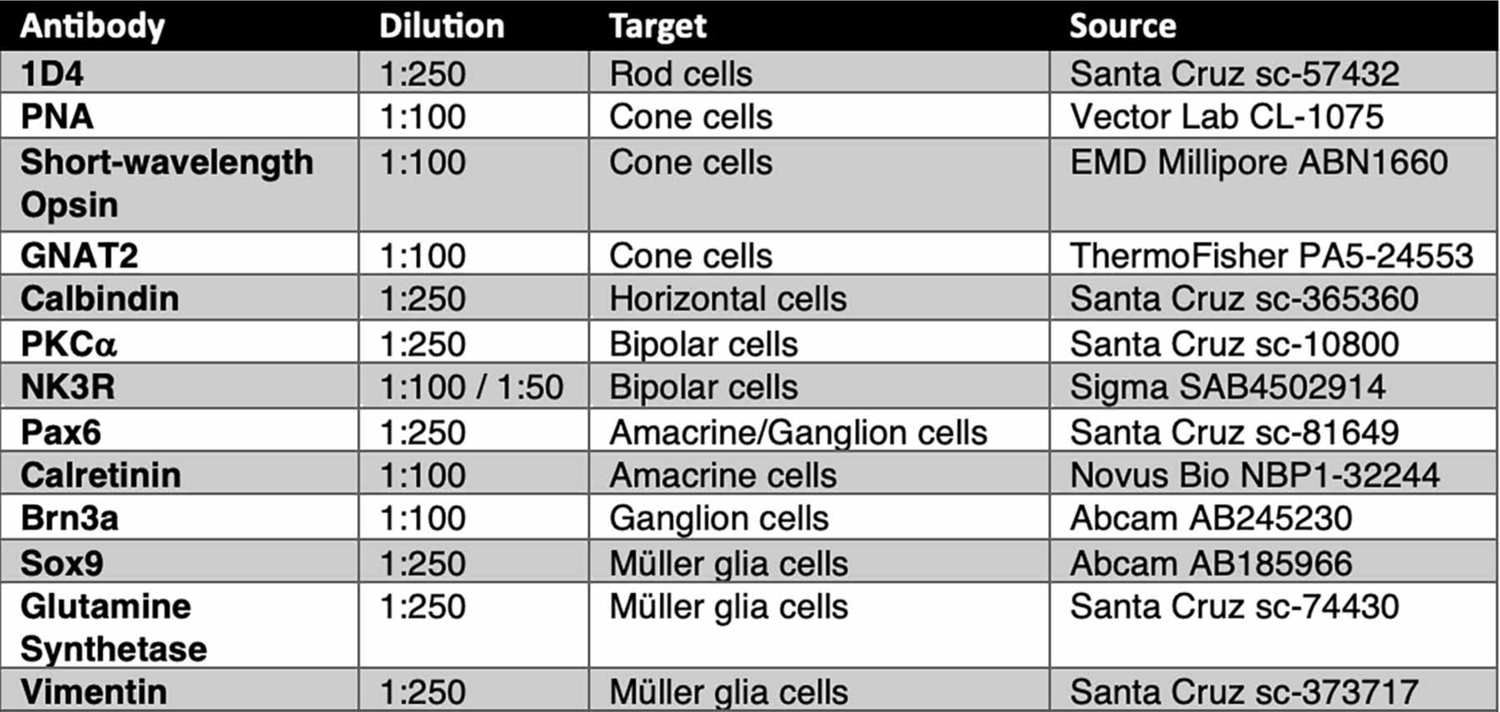
Antibodies used in this study. Antibodies used in all immunohistochemical experiments.

| **Antibody** | **Dilution** | **Target** | **Source** |
| --- | --- | --- | --- |
| **1D4** | 1:250 | Rod cells | Santa Cruz sc-57432 |
| **PNA** | 1:100 | Cone cells | Vector Lab CL-1075 |
| **Short-wavelength Opsin** | 1:100 | Cone cells | EMD Millipore ABN1660 |
| **GNAT2** | 1:100 | Cone cells | ThermoFisher PA5-24553 |
| **Calbindin** | 1:250 | Horizontal cells | Santa Cruz sc-365360 |
| **PKCα** | 1:250 | Bipolar cells | Santa Cruz sc-10800 |
| **NK3R** | 1:100 / 1:50 | Bipolar cells | Sigma SAB4502914 |
| **Pax6** | 1:250 | Amacrine/Ganglion cells | Santa Cruz sc-81649 |
| **Calretinin** | 1:100 | Amacrine cells | Novus Bio NBP1-32244 |
| **Brn3a** | 1:100 | Ganglion cells | Abcam AB245230 |
| **Sox9** | 1:250 | Müller glia cells | Abcam AB185966 |
| **Glutamine Synthetase** | 1:250 | Müller glia cells | Santa Cruz sc-74430 |
| **Vimentin** | 1:250 | Müller glia cells | Santa Cruz sc-373717 |

### Cell counting and morphology characterization

Imaging was performed on a Nikon Eclipse Ti-U inverted fluorescent microscope (Nikon Instruments, Melville, NY) or a Leica SP8 DLS confocal microscope (Leica Microsystems, Buffalo Grove, IL). To control for differences in eye size, retinal counts and measurements were taken as an average per 1000μm retinal tissue, within 3000μm of the optic nerve head. All counts and measurements were performed on retinal cryosections containing an optic nerve as a landmark. Retinal measurements and cell quantification were performed using either ImageJ (https://imagej.nih.gov/ij/) or Photoshop 6.0 software (Adobe, San Jose, CA). A replicate number of 5-7 animals was used for counts and measurements of retinal neuroanatomy. Cell counts per individual represent an average of counts taken from four, 1mm-wide tissue regions evenly spaced across the retina.

### RT-PCR and real-time quantitative RT-PCR (qPCR)

The GoScript Reverse Transcriptase System (Promega, Madison, WI) was used to synthesize the first strand cDNA from 1μg of the extracted RNA. PCR primers were designed to amplify sequences associated with opsin mRNAs (validated primer sequences are listed in **Table 2**). Faststart Essential DNA Green Master mix (Roche, Basel, Switzerland) was used to perform qPCR on a Lightcycler 96 Real-Time PCR System (Roche, Basel, Switzerland). The relative transcript abundance was normalized to Ef1α expression as the reference gene control. RT-PCR and qPCR experiments were performed with three biological replicates and two technical replicates and visualized as fold change relative to the reference gene.

**Table 2:**
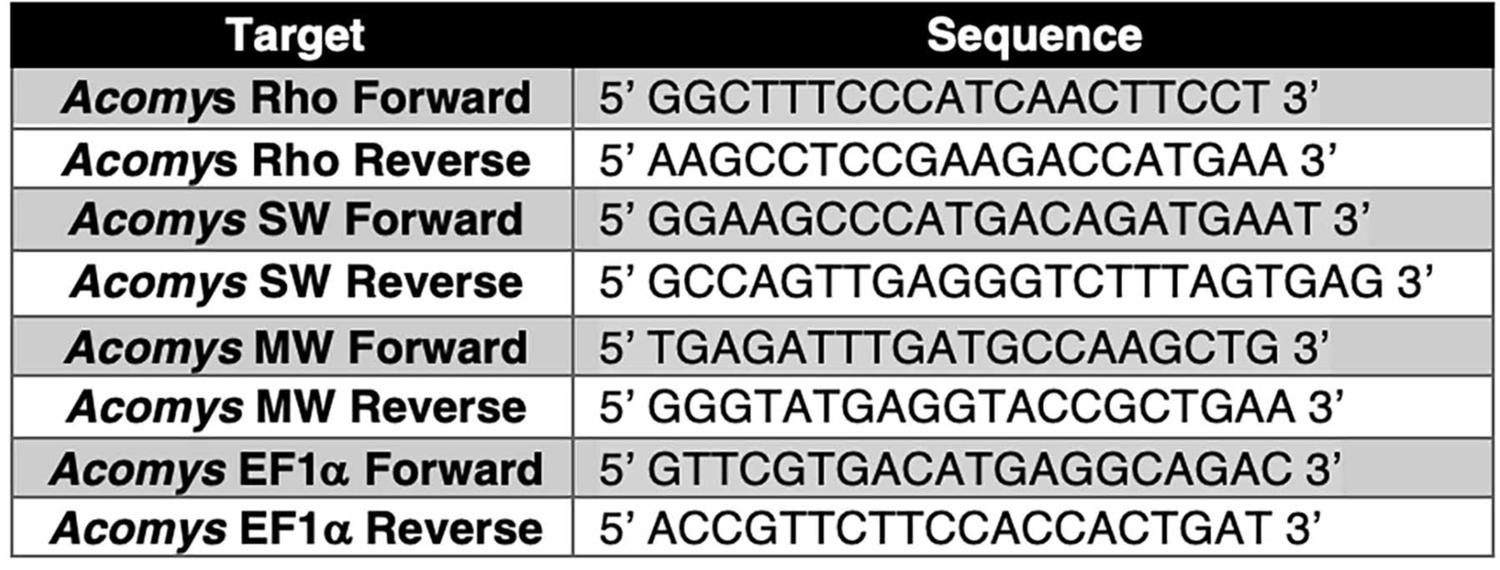
qPCR primers used in this study. Validated primer sequences used in all q-PCR experiments.

### Statistical analysis

Statistical analysis was conducted using two-factor, paired t-test and one-way ANOVA using GraphPad software. p-values less than 0.05 were considered significant and are indicated accordingly: *, p<0.05; **, p<0.01; and ***, p<0.001. All graphs were generated using Prism 9 GraphPad Software. All figures were prepared using Photoshop (Adobe version 25.1.0).

## RESULTS

### *Acomys, Meriones* and *Mus* have comparable retinal organization

Based on body length measurements (from the snout to the base of the tail), *Meriones* are the largest among the three rodent species, followed by *Acomys,* then *Mus* (Fig. 1A-A’’). Measuring the ratio of eye diameter to body length revealed that *Acomys* has a significantly larger eye per unit of body length compared to *Mus* (Fig. 1E). *Meriones* and *Acomys* eye sizes were similar to each other, with *Meriones* having a slightly larger eye than *Acomys* (data not shown). Overall, the laminar structure and layer thickness of the retina was comparable between the species, as shown by hematoxylin and eosin (H&E) staining (Fig. 1C-C’’). Measurements of the nuclear layers revealed similar widths of the INL and ONL between the species (Fig 1F, G).

**Figure 1:**
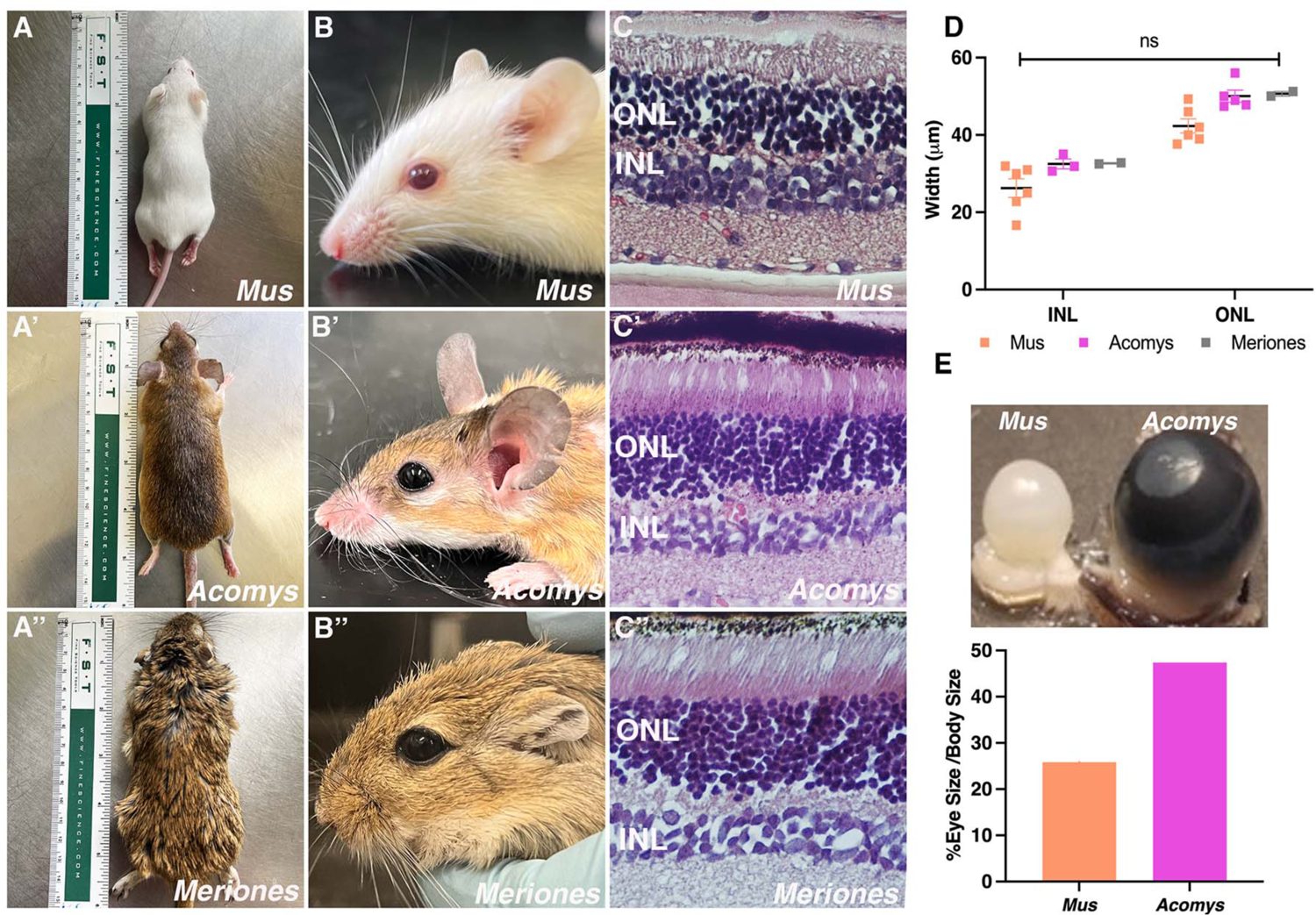
*Acomys, Meriones* and *Mus* have comparable retinal organization. (A-A’’) Body size comparisons and (B-B’’) side profile photographs of all three species (C-C’’) H&E staining of retinal cross sections. (D) Measurements of nuclear layer widths and (E) eye size comparison of *Mus* and *Acomys*. ONL, outer nuclear layer; INL, inner nuclear layer.

### *Meriones* have a higher cone density than *Acomys* or *Mus*

The extremely sensitive rod photoreceptors can detect exceptionally small quantities of light and are in high abundance in most vertebrate retinas. Immunolabeling with 1D4, an antibody that detects rhodopsin located in the rod outer segments ^41^, revealed a dense layer of rods in all three species (Fig. 2A-C’). Rods appeared long and thin, especially in *Acomys* and *Meriones* (Fig. 2B-C’). Because rods were too abundant to be individually resolved by confocal microscopy, quantification of rod density was not performed.

**Figure 2:**
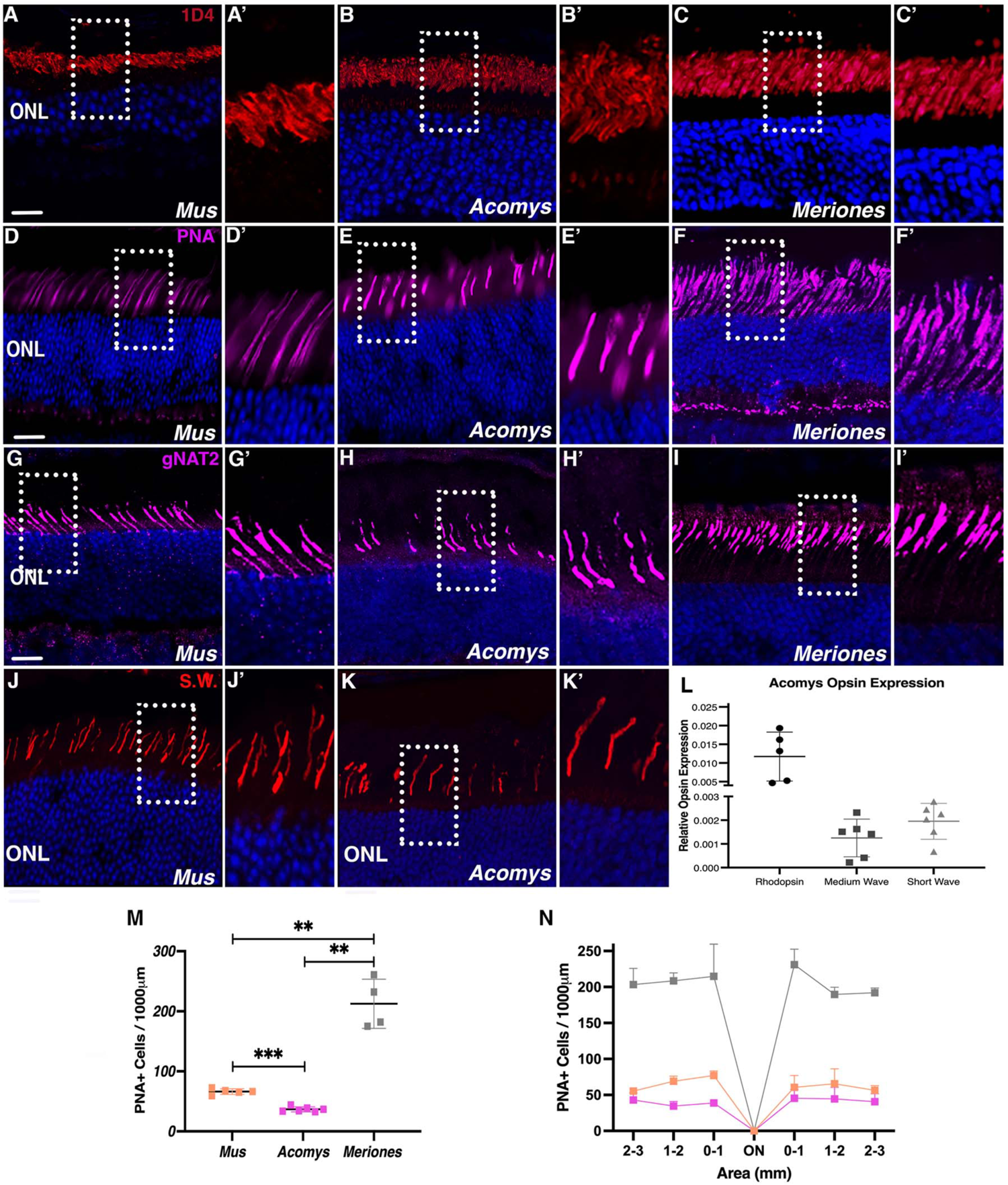
*Meriones* have an increased cone density compared to *Acomys* and *Mus*. (A-C’) Rod photoreceptor outer segments immunolabeled with 1D4 antibody. (D-F’) PNA labeling of cone outer segments. (G-I’) Cone outer segments immunostained for GNAT2. (J-K’) Short-wavelength opsin antibody labeling of cone outer segments and (L) the relative expression of opsins in *Acomys*. (M) quantification of PNA labeled cone cell density and (N) regional quantification of PNA cells across the retina. Dashed white boxes in A-K indicate the location of the enlarged region shown in the accompanying panel. ONL, outer nuclear layer. Scale bar, 20μm.

The cone photoreceptors are present in two subtypes in rodents, short-wavelength and medium-wavelength ^42^. Immunostaining for the cone marker PNA, which detects lectin on the outer segments of both cone subtypes, revealed that of the three species, *Meriones* had a significantly higher cone density than *Mus* or *Acomys*, whereas *Acomys* had the lowest cone density (Fig. 2D-F’, M). The cone outer segments in both *Mus* and *Meriones* appeared long and wispy, while in *Acomys* the outer segments appeared shorter and wider (Fig. 2D-F’). Quantifying cone abundance across regions of the retina revealed similar distributions of PNA-positive cone cells in all three species (Fig. 2N). Cones appeared in the highest densities closest to the optic nerve, with numbers declining farther away from the central retina (Fig. 2N). Similar results were obtained using an antibody for cone transducin (GNAT2; Fig. 2G-I’).

Immunohistochemistry with an antibody for short-wavelength opsin also showed that *Acomys* possess fewer short-wavelength cones per 1-mm retina than *Mus* (Fig. 2J-K). Short-wavelength and medium-wavelength opsin transcripts were able to be detected in *Acomys* using real-time quantitative PCR, and rod vs. cone transcript abundance was consistent with the immunohistochemistry results (Fig. 2L). As expected, the diurnal *Meriones* displayed the highest density of cones of the three species. In summary and somewhat surprisingly, *Acomys* (which is more active during the day than *Mus*) had the lowest density of cones of the three species analyzed.

### Bipolar cells display altered laminar positioning in *Meriones* and *Acomys* compared to *Mus*

In the mouse retina, circuit-bridging bipolar cells are present in 15 different subtypes, 14 subtypes of cone bipolar cells and 1 subtype of rod bipolar cells. Each bipolar cell subtype has a different morphology, axon length, cell body size, and cell body position in the INL ^43,44^. Using the PKCα antibody, which detects rod bipolar cells in *Mus* ^45^, we observed strong labeling of bipolar cells in all three species (Fig. 3A-C’). Distal dendrites, cell bodies and proximal terminals were all clearly distinguishable. Rod bipolar density was significantly higher in *Meriones* compared to *Mus* and *Acomys,* which were similar to each other (Fig. 3G). While cell bodies appeared mostly uniform in *Mus*, in both *Acomys* and *Meriones*, they were altered in their laminar positioning and varied in size (Fig. 3A-C, red asterisks). Classifying rod bipolar cell bodies by their location in either the proximal or distal INL highlighted the differences between *Acomys* and *Meriones* relative to *Mus* (Fig. 3B-C’, H). Both *Acomys* and *Meriones* had increased densities of bipolar cell bodies in the proximal area of INL, with *Meriones* having the highest number (Fig. 3H). Densities in the distal portion of the INL showed no difference across species (Fig. 3H).

**Figure 3:**
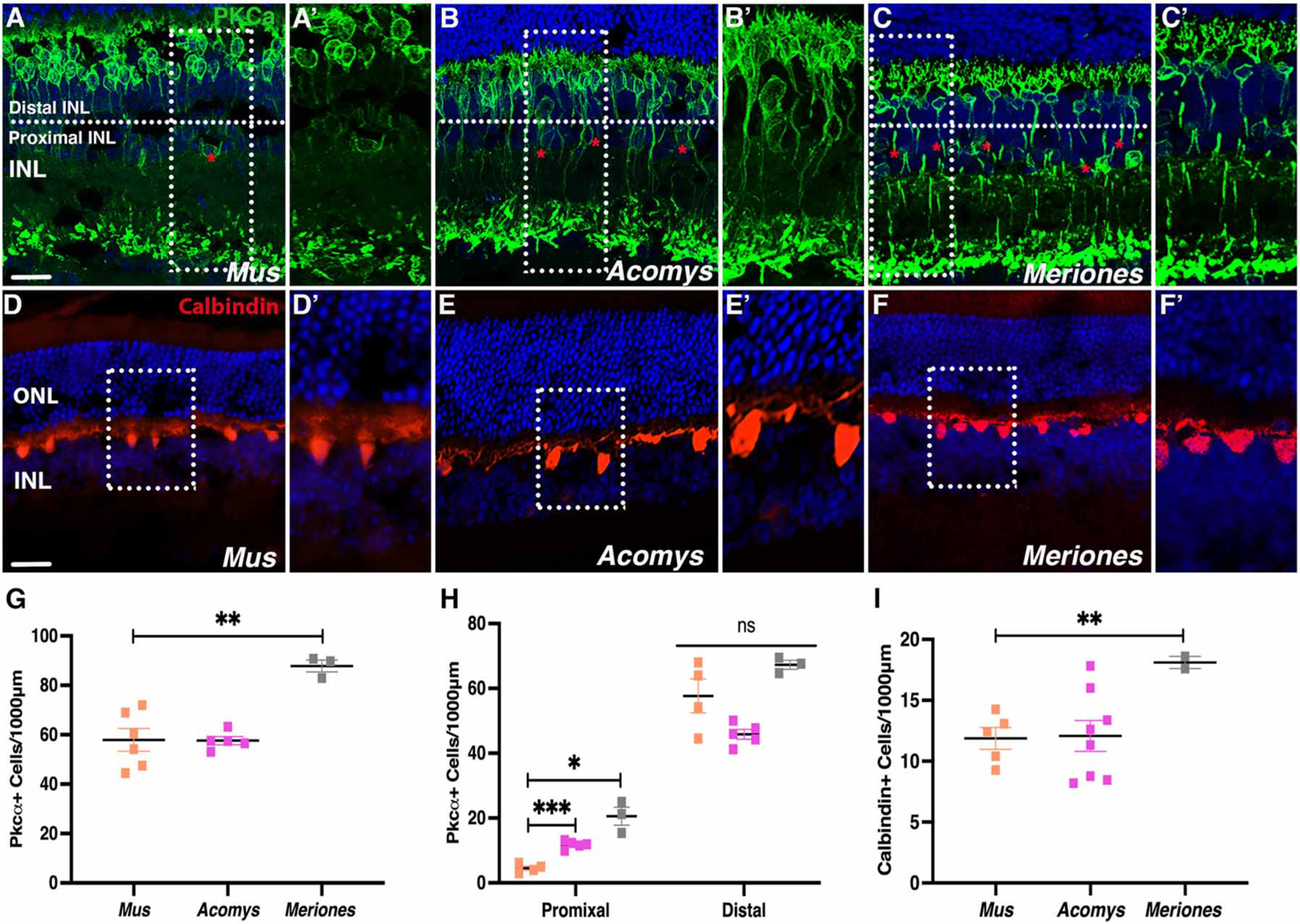
Bipolar cells have increased densities and altered laminar positioning in *Meriones* and *Acomys* compared to *Mus.* (A-C’) Rod bipolar cells immunolabeled with PKCα. (D-F’) Horizonal cells visualized with antibody against calbindin. (G) Quantification of total and (H) region densities of bipolar cells (regional differences indicated by red asterisks). (I) Quantification of calbindin labeled horizontal cells. Dashed white boxes in A-F indicate the location of enlarged region shown in the accompanying panel. ONL, outer nuclear layer; INL, inner nuclear layer. Scale bar, 20μm.

Retinal horizontal cells provide feedback and feedforward information to the photoreceptors and aid in contrast and color correction ^46^. The mouse retina contains one horizonal cell subtype ^47^. Using an antibody against calbindin, the calcium-binding subunit of horizontal cells, horizontal cell bodies and axons were visible in all three species and were especially clear in *Acomys* (Fig. 3D-F’). When quantified, horizonal cell density was higher in *Meriones* compared to both *Mus* and *Acomys* (Fig. 3I), although *Acomys* showed quite a bit of variation in the samples analyzed.

### Amacrine cells have increased density in *Acomys* and *Meriones* compared to Mus

Amacrine cells are present in approximately 30 subtypes across mammals ^48^. Using a Pax6 antibody, which detects all retinal amacrine cells and subsets of ganglion cells in *Mus* ^49^, we detected an overall similar distribution in *Acomys, Meriones*, and *Mus*; quantification of Pax6-positive cell density in the INL showed no significant difference between the three species (Fig. 4A-C’, G). However, using the amacrine-specific marker calretinin, which labels the calcium-binding protein in starburst amacrine subtypes in mice ^50^, the density of this amacrine cell subtype appeared greater in *Acomys* and *Meriones* compared to *Mus* (Fig. 4D-F’). Quantification of calretinin-positive cells confirmed that both *Acomys* and *Meriones* had an overall significantly higher density of calretinin-positive amacrine cells than *Mus*, and the distribution of these cells across regions of the retina was similar between the two species (Fig. 4H-I).

**Figure 4:**
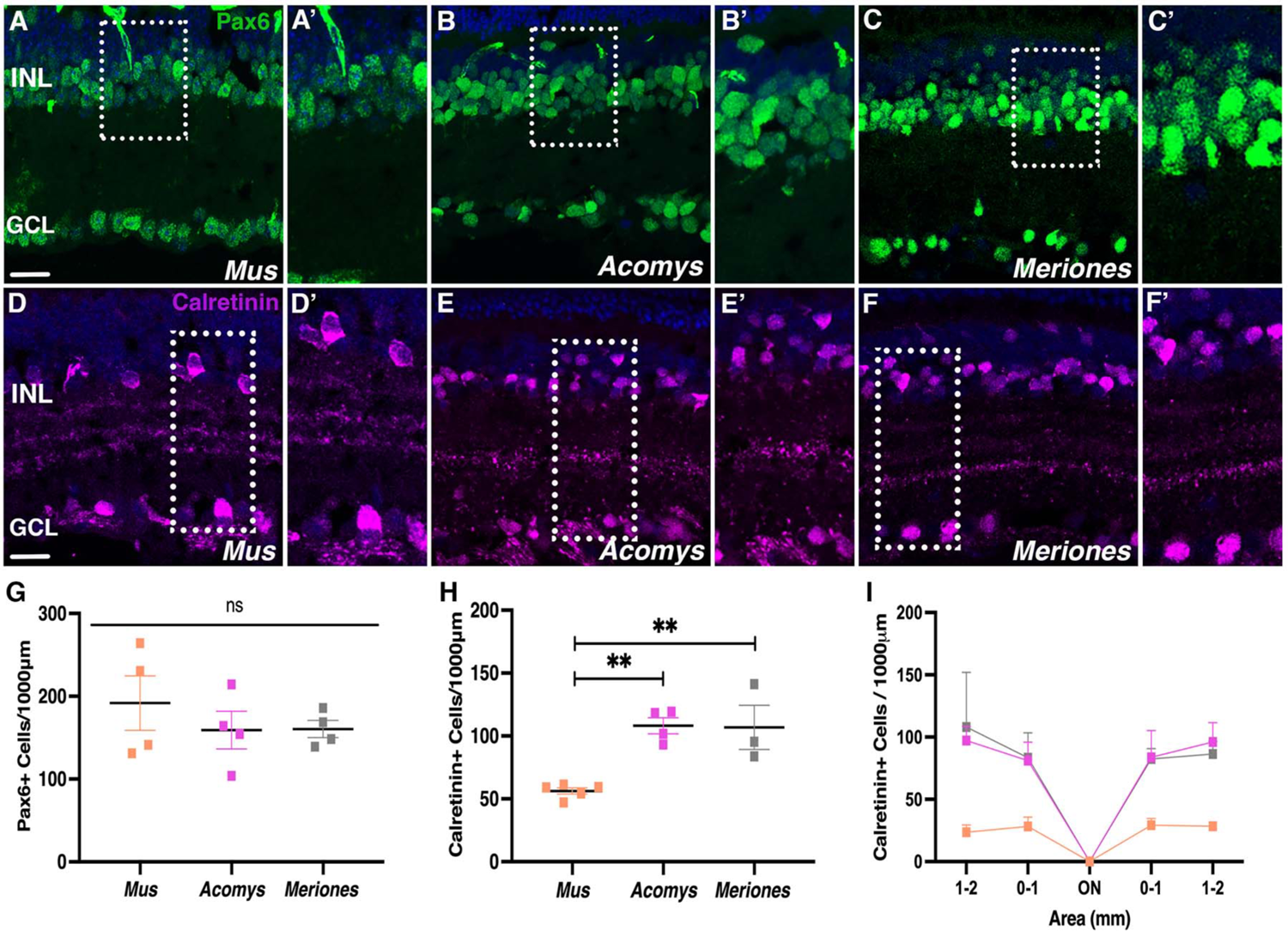
Amacrine cells have increased density in *Acomys* and *Meriones* compared to *Mus.* (A-C’) Immunostained Pax6-positive amacrine cells. (D-F’) Subtype specific amacrine cells immunolabeled with calretinin. (G) Quantification of Pax6-positive cell density. (H) Quantification of calretinin-positive cells and (I) regional distribution of calretinin cells across the retina. Dashed white boxes indicate in A-F indicate the location of the enlarged region shown in the accompanying panel. ONL, outer nuclear layer; INL, inner nuclear layer; GCL, ganglion cell layer. Scale bar, 20μm.

The retinal ganglion cells, the output neurons of the retina, are a diverse class of neurons with over 30 subtypes ^51^. Immunolabeling for Pax6-positive cells in the GCL revealed similar ganglion cell density and position between all three species (Fig. 5A-C’, G). Immunostaining for Brn3a, which is specific for vision-forming retinal ganglion cells ^52^, indicated an increase in density in *Acomys* compared to *Meriones* and *Mus* (Fig. 5D-F’, H). Quantification confirmed similar distributions of ganglion cells across the retina in *Acomys, Meriones*, and *Mus* (Fig. 5D-F’, H-I).

**Figure 5:**
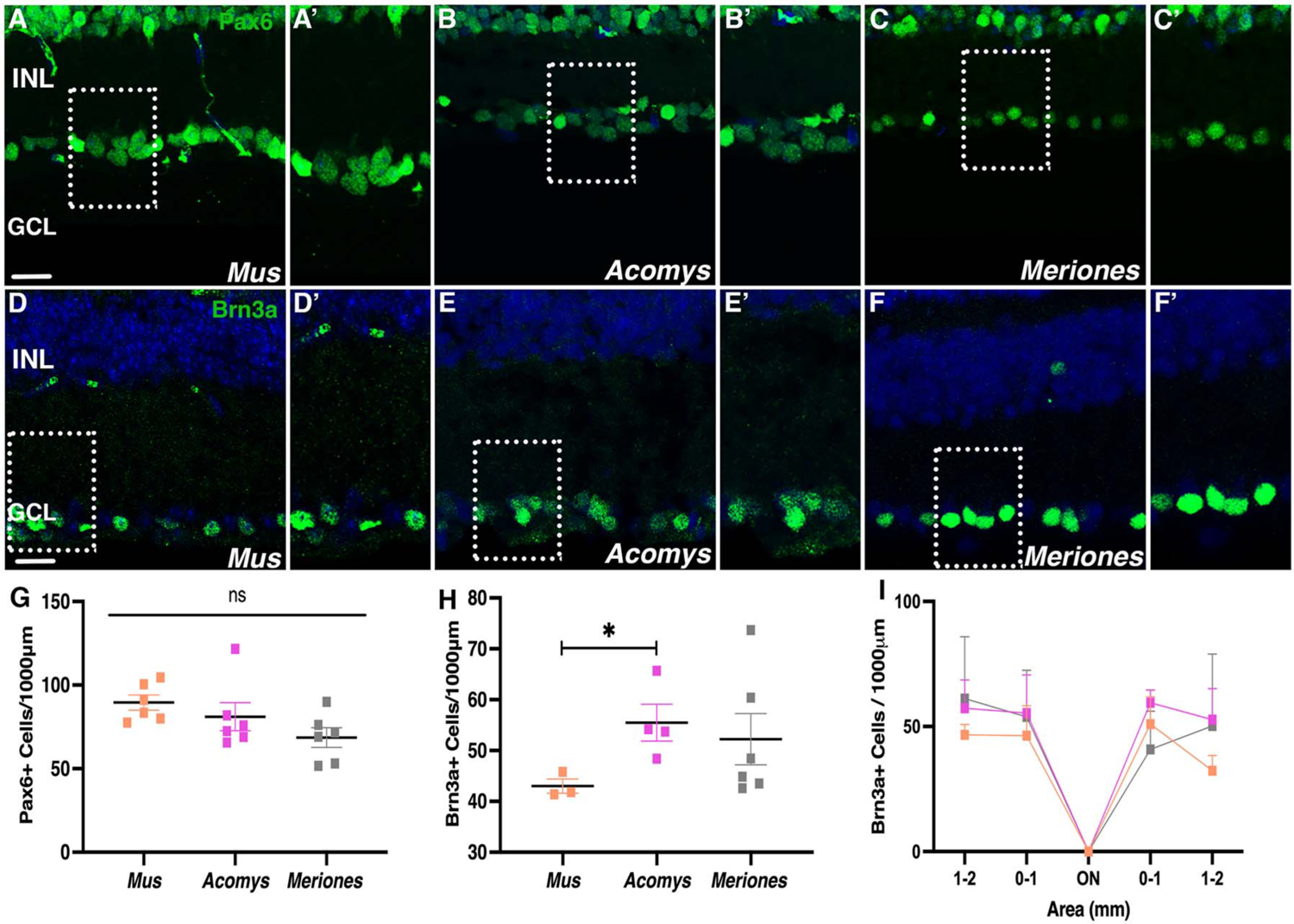
Ganglion cell density is comparable between *Acomys, Meriones* and Mus. (A-C’) Retinal ganglion cells immunolabeled with Pax6, (D-F’) Antibody staining against ganglion cell subtype specific marker Brn3a. (G) Quantification of Pax6 labeled cell densities. (H) Quantification of Brn3a positive cells and (I) regional distribution of Brn3a cells across the retina. Dashed white boxes in A-F indicate the location of the enlarged region shown in the accompanying panel. INL, inner nuclear layer; GCL, ganglion cell layer. Scale bar, 20μm.

### Müller glia display similar morphology and density in all three species

Finally, we analyzed the morphology and distribution of the Müller glia, the resident glial cell of the retina, which are critical for retinal homeostasis, structural support, and mediate regeneration in some species ^53^. Antibody labeling against glutamine synthetase revealed the long Müller glial processes which span the entire width of the retina ^54^. Müller glial morphology appeared similar in all three species; however, the cell body positioning appeared less uniform in *Acomys*, along with uneven positioning of the Müller end-feet (Fig. 6A-B’). *Meriones* Müller cells had a lower affinity for the glutamine synthetase antibody and were not visualized as easily (Fig. 6C-C’). Immunostaining with vimentin, an intermediate filament protein found in the cytoskeleton of Müller cells, highlighted the end feet of the Müller glia and their long and thin proximal processes in all three species (Fig. 6D-F’). Immunolabeling for the transcription factor Sox9, which in the adult retina is specifically expressed in Müller cell bodies, revealed *Acomys* and *Mus* to have similar densities, while *Meriones* showed a higher density of Müller glia in the INL (Fig 6G-I’, J). Interestingly, whereas the Müller cell bodies in *Mus* and *Meriones* displayed a consistent polygonal shape, *Acomys* Müller cell body shape was much more variable (Fig. 6H-H’). Overall, the density of Sox9-positive cell bodies across the four retina regions was similar between all three species (Fig. 6K).

**Figure 6:**
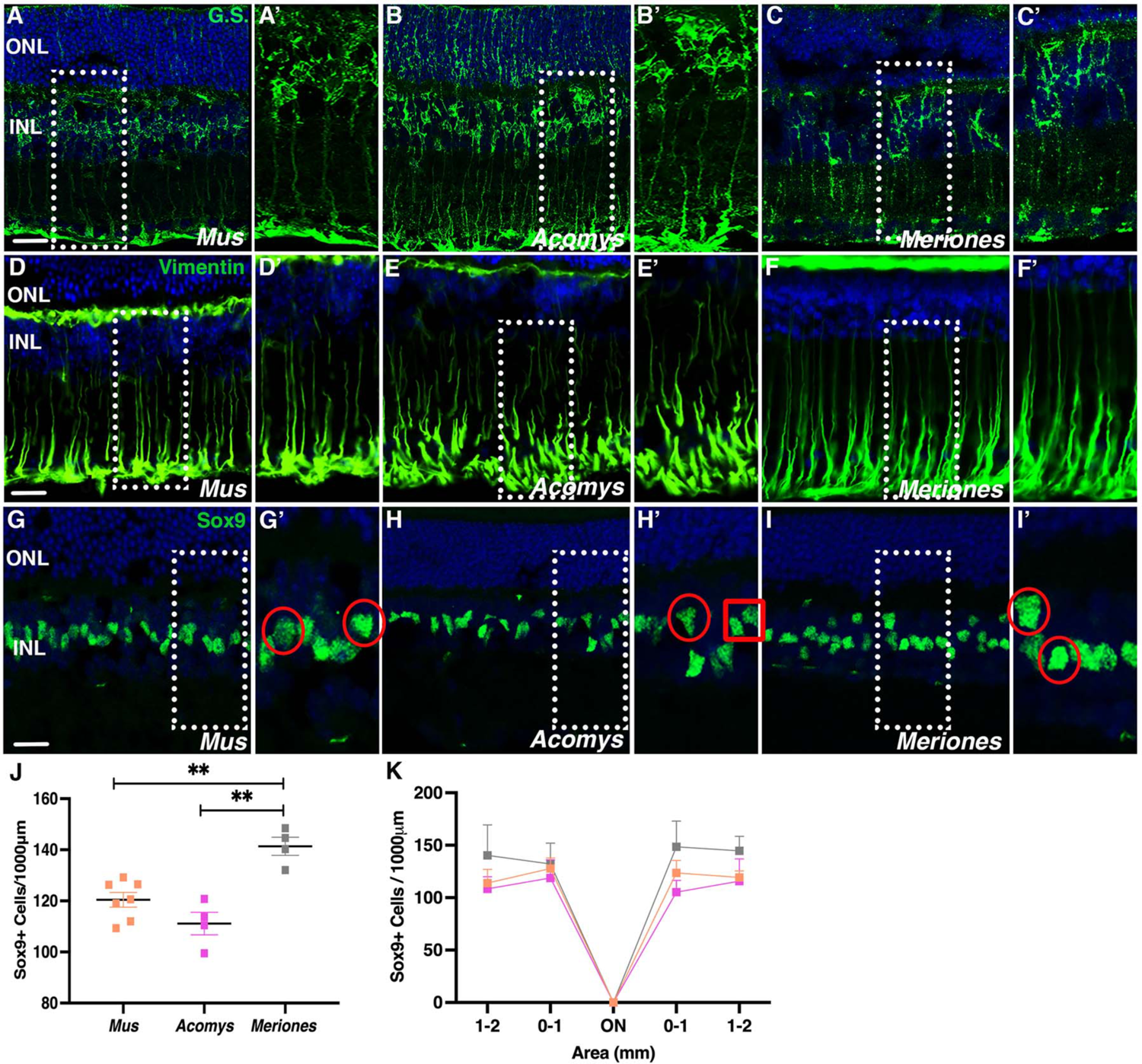
Müller cells display similar morphology and density across all three species. (A-C’) Müller glia cells immunolabeled with glutamine synthetase. (D-F’) Antibody against vimentin labeled Müller cells. (G-I’) Sox9 immunolabeling of Müller cell bodies, (red circles indicate examples of differences in cell shape). (J) Quantification of Sox9 labeled cells and (K) regional distribution of Sox9 labeled cells across the retina. Dashed white boxes in A-I indicate location of the enlarged region shown in the accompanying panel. ONL, outer nuclear layer; INL, inner nuclear layer; G.S., glutamine synthetase. Scale bar, 20μm.

## DISCUSSION

Here we established the basic cellular and molecular anatomy of the retina in *Acomys dimidiatus* and *Meriones unguiculatus*. Our detailed characterization now provides a platform to use these animals as new mammalian models for retinal cell biology and regeneration. Introducing more diversity of animal models for vision biology is crucial to expanding our understanding of the range of visual capabilities and adaptations across the animal kingdom. Furthermore, retinal degenerative eye diseases such as macular degeneration and retinitis pigmentosa affect millions of people worldwide ^55,56^. Studying the retinas of animals with good daytime vision (such as *Meriones*) or enhanced regenerative abilities (such as *Acomys*) could facilitate discoveries that open new therapeutic avenues to treat these sight threatening conditions.

The genus *Meriones* has a rich scientific history in research areas such as reproduction, diabetes, epilepsy, and aging ^12–14^, but published investigations using this species had declined until recently. *Meriones* are an uncommon diurnal rodent with similar visual function as humans ^16^. Using ERG analysis and immunocytochemistry with opsin antibodies, previous reports described a high density of medium and short-wavelength sensitive cones in the *Meriones* retina ^20^. Our observations with PNA staining and GNAT2 antibody labeling (**Fig. 2**) confirm and extend this result, which correlates well with the diurnal pattern of activity in this species.

A previous study demonstrating functional recovery from complete spinal cord transection in *Acomys* raised the exciting prospect that these animals have the capacity for enhanced central nervous system regeneration ^35,57^. Investigations of retinal regenerative capacity in *Acomys* will require an understanding of their retinal anatomy as well as a panel of reliable retinal cell type markers, and so the analysis provided here is the first step in this endeavor. Considering *Acomys* are not strictly nocturnal, and given the reported flexibility in their sleeping patterns ^40^, we hypothesized they would possess a higher density of cone photoreceptors compared to *Mus.* However, we observed the opposite result; immunolabeling for GNAT2 and short-wavelength opsin, as well as PNA staining, revealed that *Acomys* have the lowest density of cones of all three species in our study. (**Fig. 2**). It is possible that *Acomys* possess additional cone subtypes that weren’t detected by our methods; however, this seems unlikely as there were no additional cone opsin genes identified in the *Acomys* genome (EMBL: https://www.ebi.ac.uk/ena/browser/view/GCA_907164435.1). Furthermore, the long and thin rod photoreceptor morphology we observed in *Acomys* has been previously associated with animals with more dark-adapted vision ^58^. There is currently no published work examining retinal function in *Acomys*, either through behavioral assays or electroretinography. To determine if *Acomys* have any visual adaptations that support increased levels of daytime activity, additional experiments such as extensive ERG analysis and behavioral observations need to be completed.

Although both *Acomys* and *Meriones* retinas were comparable to *Mus* in basic structural organization and cell type composition, some distinct differences were observed. For example, we observed altered laminar positioning of some bipolar cell bodies (compared to *Mus*) labeled with the PKCα antibody, which marks rod bipolar cells in *Mus* (**Fig. 3**). In both species, but especially *Meriones,* we observed more cell bodies located in the proximal region of the INL than in *Mus*. Given that the location of the cell body has been correlated with the bipolar cell subtype ^59,60^, one explanation for our results could be that PKCα detects additional subtypes of bipolar cells in *Meriones* and *Acomys* to those labeled in *Mus.* Although traditionally, PKCα labels mainly rod bipolar cells in mammals, not enough diurnal rodents, or ones with flexible sleeping patterns, have been examined in order to rule out its expression in additional subtypes^61^. Alternatively, PKCα could be labeling other types of retinal neurons in addition to bipolar cells in *Acomys* and *Meriones*. For example, some studies have reported that PKCα can label some amacrine cells ^61,62^. We attempted to visualize other bipolar cell subtypes using an antibody to NK3R, which has been shown to label two different types of ON bipolar cell in the mouse ^63^. However, the NK3R antibody did not have sufficient cross-reactivity with *Acomys* or *Meriones* retinal cells (data not shown). Additional markers for bipolar subtypes need to be established to examine the full complement of bipolar cells across these species.

There are over 30 subtypes of amacrine cells and 15 subtypes of ganglion cells in mammals ^44,64,65^. We found that both *Meriones* and *Acomys* had increased densities of calretinin-positive amacrine cells compared to *Mus*, yet similar numbers between each other (**Fig. 4**). Other amacrine subtypes, however, could have more similar abundance between the species since we did not detect significant differences in Pax6-positive cell densities in the INL. The increase in calretinin-positive amacrine cells (relative to *Mus*) was obvious at first glance. The higher number of amacrine cells in *Meriones* could be related to the increased densities in both their cone and bipolar cells relative to *Mus*, which may necessitate additional circuit feedback. It is less clear why *Acomys* had a higher density of amacrine cells than *Mus*. Given that amacrine cells play a crucial role in in the perception of vision in starlight, twilight, and daylight ^66^, perhaps the increased amacrine cell number is related to their distinctly different movement patterns throughout the day than *Mus,* which could necessitate the need for increased amacrine processing.

In conclusion, this study establishes a suite of commercially available antibodies that can be used to detect retinal cell types in *Meriones* and *Acomys*, two rodent models with unique qualities that make them valuable for understanding the diversity of mammalian visual behavior and retinal cell biology. Future work will focus on analyzing visual function and response to retinal injury in these models.

## ACKNOWLEDGMENTS

We would like to thank Ben Heckman for help optimizing various antibodies. We thank Josh Sarli and other members of the Seifert lab for exceptional animal care. We also thank members of the Morris lab for valuable input and technical assistance. This work was supported by grants from the Retina Research Foundation (to A.C.M. and A.W.S); J. Bills was supported by a NSF GRF.

